# *Tetragonisca angustula* resin gathering behaviour of *Schizolobium parahyba*

**DOI:** 10.1101/2021.09.09.459628

**Authors:** Leonardo Regnier

**Affiliations:** University of Sao Paulo

**Keywords:** Propolis, Stingless bees, Resin Foraging, Meliponini, Physiology

## Abstract

*Tetragonisca angustula* is a very important stingless bees species. This study aimed to report the resin gathering behaviour of this species of a possible new resin source: *Schizolobium parahyba*. Evaluations of temperature, number of bees in gathering activity, time and season were used to characterize bee’s behaviour. Resin gathering activities were concentered between 10 and 14 hours, with a moderate linear positive correlation index with the temperature. No bee activity was observed when temperatures were below 16.69°C. Gathering suffered extreme reduction during winter and greater activity in summer. Bees exploration were concentered on the younger leafs and apical portion of *S. parahyba*. Older leafs were mainly ignored, and gradually the exploring activity was constantly migrating to most young parts, while apical exploration was consistent all the studied period.

## Introduction

*Tetragonisca angustula* is a native stingless bee from the Pantropical region. Belonging to the Meliponini tribe, this species compose one of the most diverse groups of bees (Divya et al. 2020). This group is one of the most important to the tropical region, due to its ecological importance of pollination to all the angiosperms of this part of the world (Sawaya et al. 2009).

Recently this group of bees have been explored as an alternative in controlled crop pollination, and greenhouse cultures (Borges and Blochtein 2005) offering almost no harm to humans, since they lack the usual sting present on other bee groups, they are better alternatives (Da Silva et al. 2011).

Stingless bees gather plant resins to build nest structures and use them in the composition of propolis to protect the hive (Grüeter 2020). This is mainly attributed to the higher moisture content inside the nest, which could spoil provisions if pots are made purely of wax (Melo 2020). However, different from what happens for the collection of pollen and nectar, which are easily studied through observational studies, the recognition of the sources of resins used by bees to compose the propolis and nest structure is scarce (Akatsu and Soares 2009).

In general, direct observation of resin foraging behaviour is extremely difficult, especially because in bushlands this activity can be sparse and occur in the canopy of trees (Bankova et al. 2019). Therefore, most of the studies compare chemical composition to indicate the propolis origin (Sawaya et al. 2009). Besides that, the identification of resin sources is important, since propolis is a commercially relevant product from beekeeping (Bankova, Popova, and Trusheva 2018).

*Schizolobium parahyba* (Vell.) Blake is a native tree species of Brazil (Romão and Mansano 2020). Present on several biomes is a pioneer species (Campião, Garlet, and Evangelista 2021). Due to its fast growth and resistance to environmental conditions, it has been presenting very promising economical uses such as reforestation and wood industry (Campião, Garlet, and Evangelista 2021).

Thus, given the specific conditions in which the direct observation of the resin gathering was possible. This study aimed to report the activity of bees collecting resins from direct observation of trees.

## Material & Methods

This study was conducted in Cotia-Brazil (23°35’55.7”S 46°50’10.5”W). The experiment was conducted between February and June 2021 consisting of the observation of foraging bees on a young individual of *Schizolobium parahyba* (being possible to observe the complete individual). Leafs were numbered from basis to apex, and the number of bees exploring each plant part counted during 1 chronometer minute.

The data comprehended 302 observations during the studied period of each 15 minutes from 6:00 am and 7:00 pm. The shade air temperature was measured using a mercury thermometer.

Data analysis was conducted using R (R Core Team 2020). Data manipulation consisted of the use of tidyr (Wickham 2020), and plots were performed using the ggpubr (Kassambara 2020) package.

## Results & Discussion

### Propolis gathering behaviour

Several bees explore the same individuals of *S. parahyba* (Figure 1A). First of all, the field bees patrol the leafs, probably looking for clues of the most profitable resin sources. After choosing the plant part, and landing, the worker bee uses its mandibles to scrape the plant surface to remove the resin (Figure 1B). A very similar process to the described to *Macaranga thanarius* exploration in the production of Okinawan propolis (Kumazawa et al. 2008).

**Figure 1:**
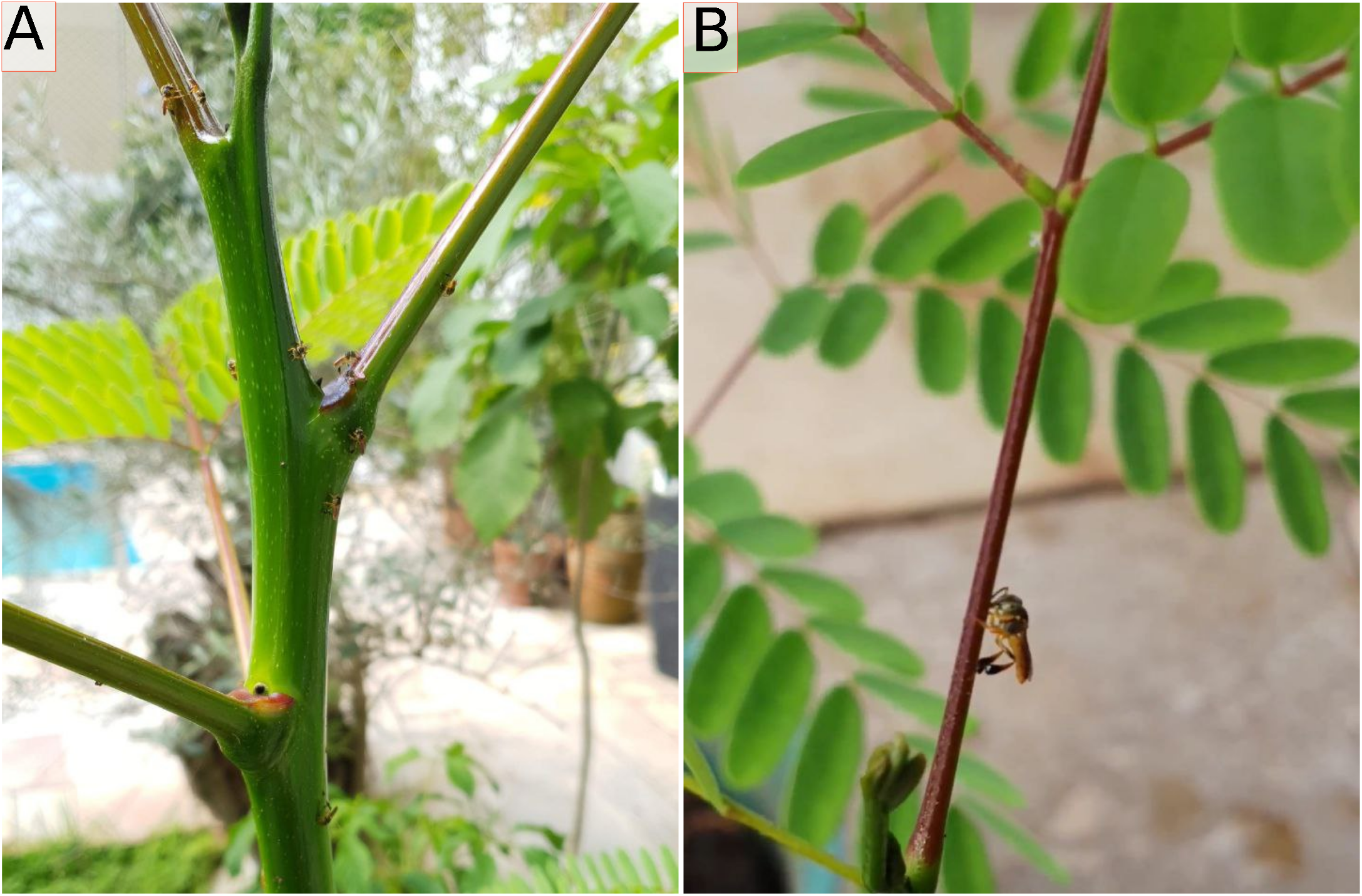
Worker bees gathering resins from young parts from individuals of *S. parahyba*. **A**- Several individuals exploring the apical portion of the stem. **B** - Young leaf resin gathering.

Gradually the bees form pellets of resin in the mouth and transfer it to corbiculae. Usually is mentioned that when stingless bees need to break off pieces of the resin exudates, the whole process would take between 7 min and 1 hour depending on the weather condition (Çelemli 2013).

During the studied period, maybe because the workers just need to abrade the surface of plants, the process used to be a little faster than 7 minutes, but this was also affected by the environmental conditions.

As a whole, the petiole and pulvinus were the parts bees seems to prefer to explore. This is expected since the secretion of resins is very variable between the leaf parts in Fabaceae (Langenheim et al. 1982). Other species of Meliponini were also observed exploring *S. parahyba*, but this study focused only on *T. angustula*.

### Gathering activity

Comparing the number of bees dedicated to the gathering activity, the resin collection seems to peak between 10-14 hours, with a gradual rise and reduction over the day (Figure 2). Very similar to what was previously described as general hive activity, which peaks between 11 and 13 hours (Kleinert-Giovannini and Imperatriz-Fonseca 1986).

**Figure 2:**
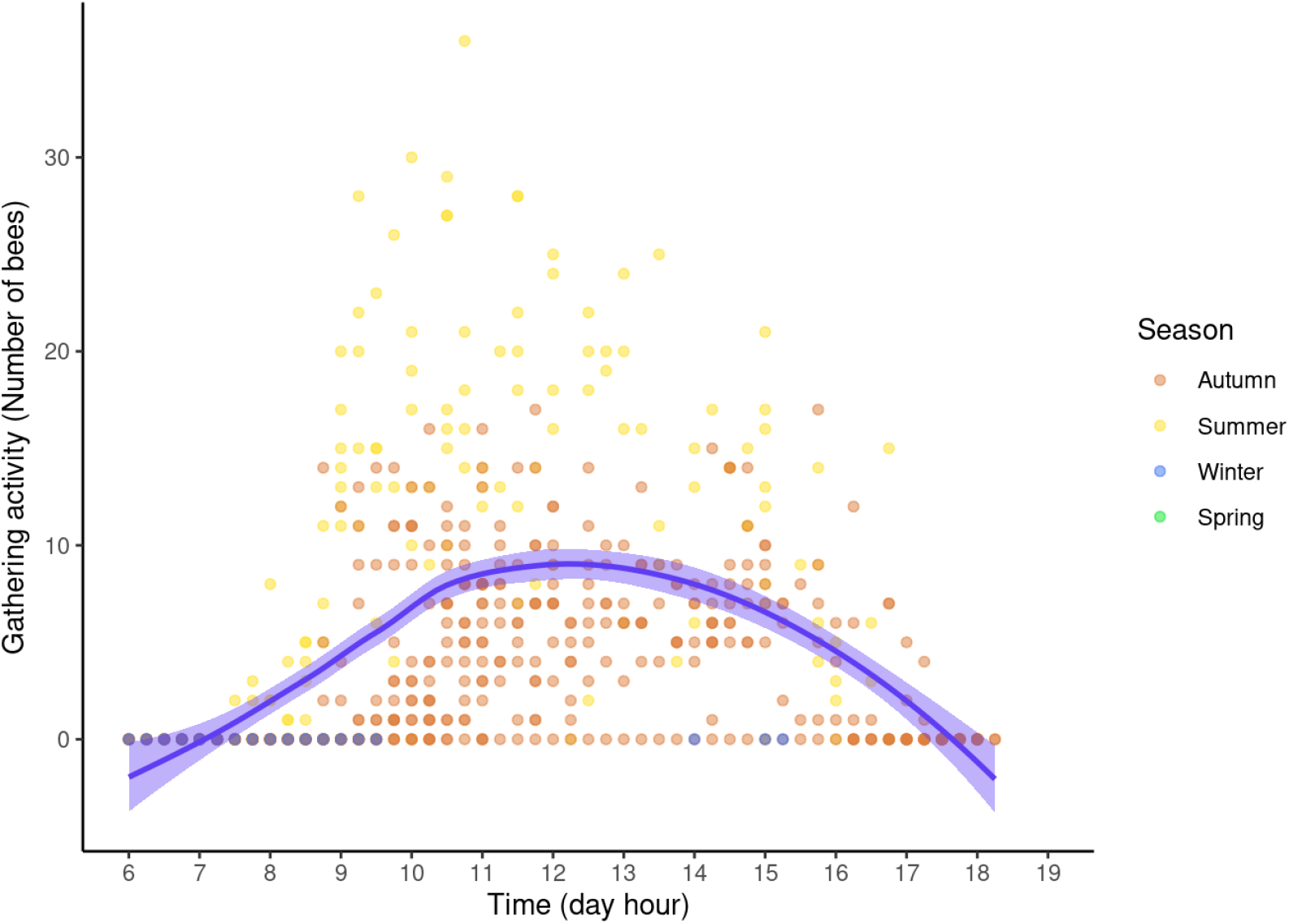
Number of bees in resin gathering activity during day hours.

In general, the end of foraging activity is mainly determined by light decrease, rather than temperature reduction (Figueiredo-Mecca, Bego, and Nascimento 2013). This is coherent with the obtained data, since the temperature usually present a constant pattern at the end of the day, starting to reduce only after the sunset (data not shown).

Flight activity is described to start with only a few members of the hive, and gradually increase with temperature and sunlight raising (Figueiredo-Mecca, Bego, and Nascimento 2013). This gradual increase was also obtained in resin gathering activity (Figure 2). The cessation of the activity is mainly attributed to the end of sunlight, due to the fact that even with temperatures being 1 until 4 °C higher than the minimum temperature required for external activity, bees stop the foraging behaviour (Kleinert-Giovannini and Imperatriz-Fonseca 1986).

Other studies mention that external activity is mainly affected by sunlight, temperature, air humidity, and wind (Figueiredo-Mecca, Bego, and Nascimento 2013). Besides that, in pollen gathering, the activity is mostly concentrated during the morning, which is attributed to the restricted availability of this resource (Bisui, Layek, and Karmakar 2019). In contrast to this pattern, resin gathering seems not to be affected. This is somehow expected since the resin collection does not depend on floral resources, which feasibility of exploration varies between day hours.

### Temperature and gathering behaviour

Temperature seems to be intermediate positively correlated with the resin gathering (Figure 3), presenting a moderate correlation index (Dancey and Reidy 2020). From the obtained linear model, it is possible to infer that at temperatures below 16.69 °C there is no resin gathering activity. This is a very close result to the mentioned minimum temperature to the external activity found in *Scaptotrigona depilis* (Figueiredo-Mecca, Bego, and Nascimento 2013). Nevertheless, is even closer to the minimum temperature to external activity documented to *Melipona* genus, which comprehends temperatures between 16 and 17 °C (Kleinert-Giovannini and Imperatriz-Fonseca 1986).

**Figure 3:**
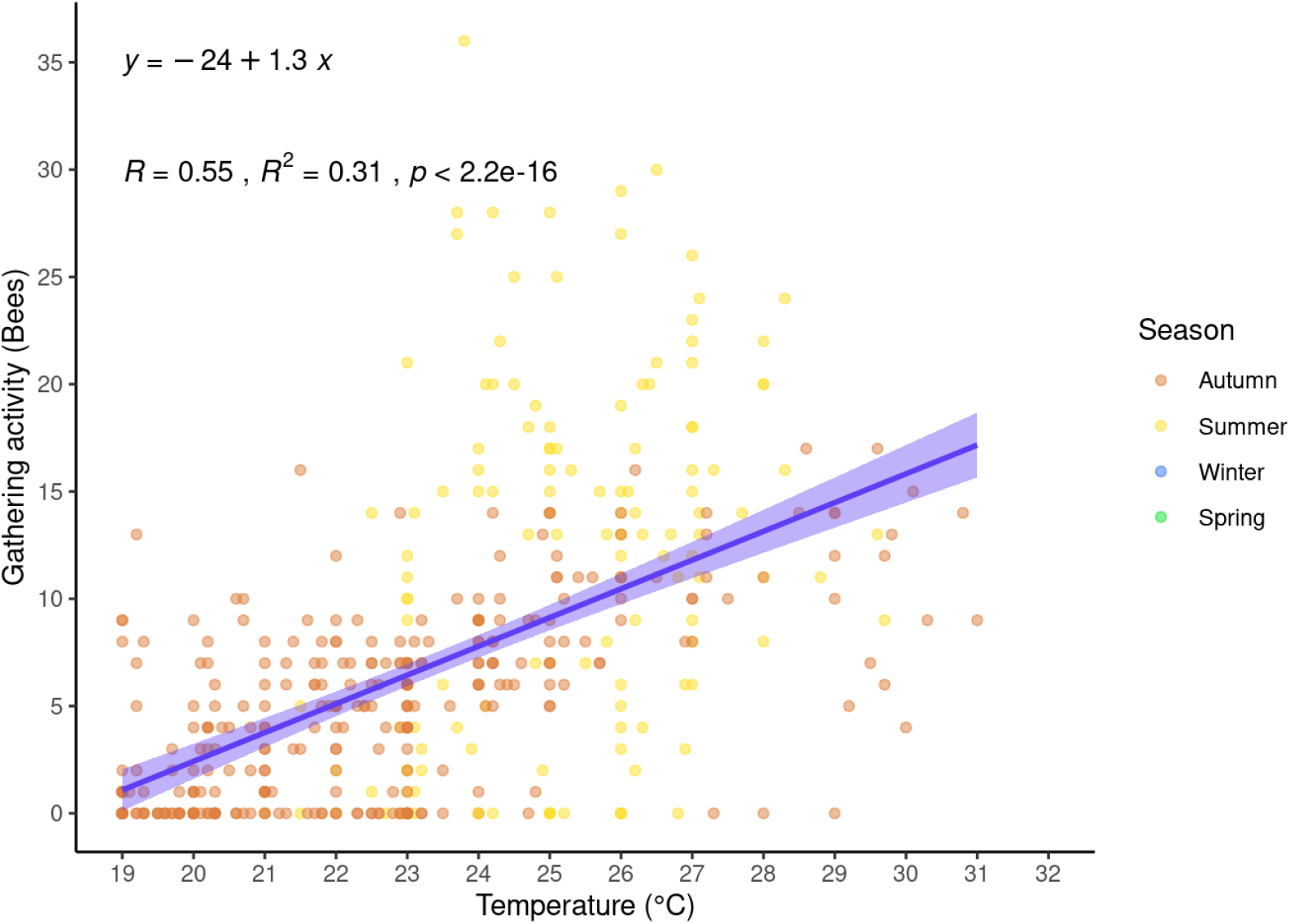
Scatter plot of temperature and gathering activity of bees.

### Gathering between seasons

Other studies mention that external activity is mainly affected by sunlight, temperature, air humidity, and wind (Kleinert-Giovannini and Imperatriz-Fonseca 1986). However, their degree of influence is not the same.

Effect of seasons in the resin gathering was observed. The warmer months have concentrated most of the gathering activity (Figure 4), in which summer had the greater number of worker bees gathering resins, autumn have a gradual reduction, while no activity was seen in winter.

**Figure 4:**
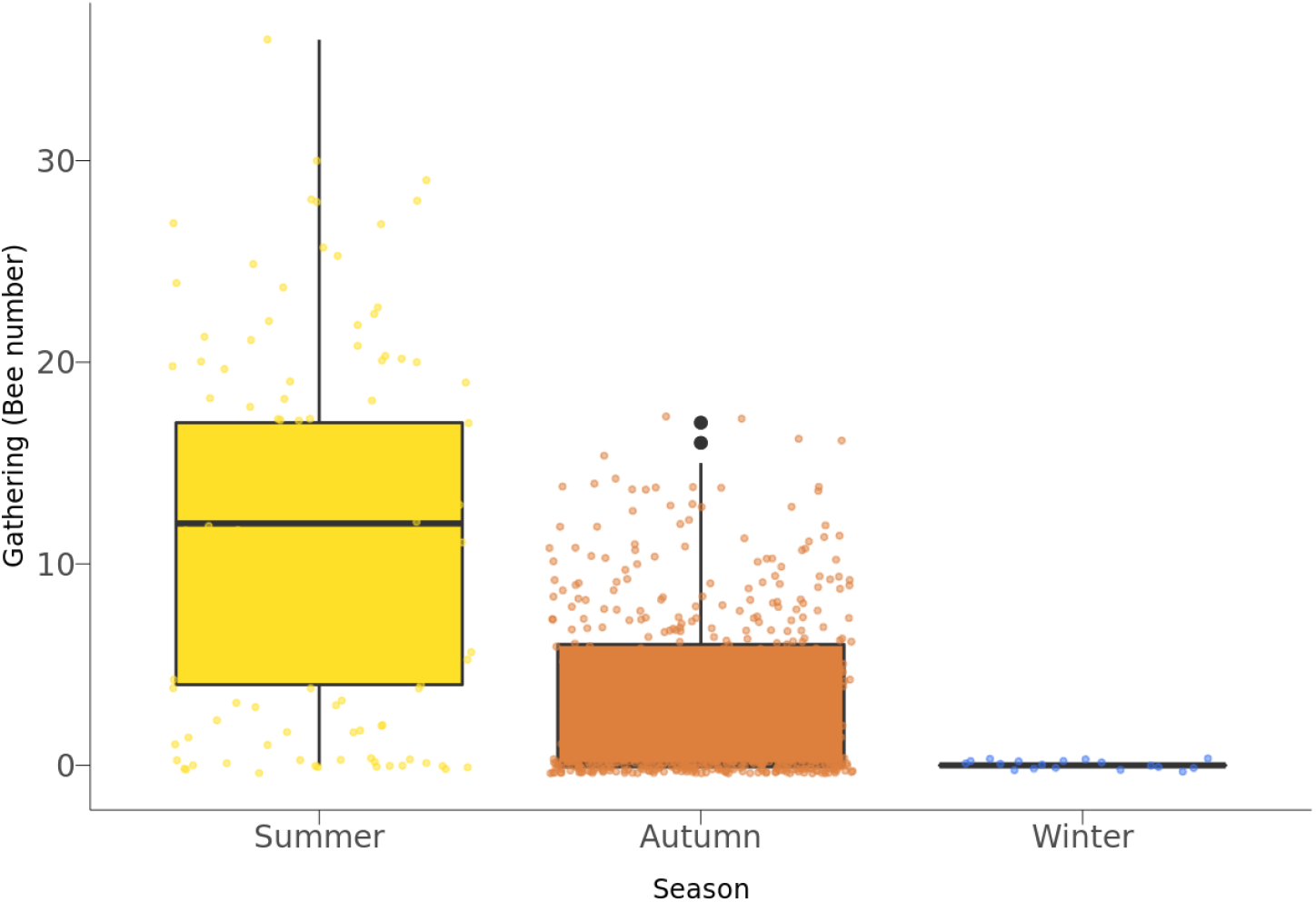
Number of bees in resin gathering activity during day hours.

Resin gathering is mentioned to be more intense in late summer and autumn due to the availability of flower resources, which is more scarce in this period (Bankova, Popova, and Trusheva 2018). The present data corroborates this information, considering that the spring activity was not contemplated, and future studies are required to properly attest to this hypothesis.

Despite of this study biases, the obtained information is accordingly to the typical seasonal pattern found in the Southeast of Brazil, which consisted of the external activity increased in the wet season in warm months (August until March) and decreased during the cold and driest months (April until July) (Figueiredo-Mecca, Bego, and Nascimento 2013). Directly impacting the resources gathering.

Fabaceae members seem to be responsible for most of the pollen load of some stingless bees species (Bisui, Layek, and Karmakar 2019). As well as have been indicated as possible resins sources, because the presence of isoflavones, an important chemical marker of this family, has been found on several propolis types (Bankova, Popova, and Trusheva 2018).

Besides that, *Hymenaea courbaril*, a member of Fabaceae, have already been indicated as one of the resin sources to *S. depilis* propolis (Akatsu and Soares 2009).

### Tree exploration

*S. parahyba* individuals has a sticky material covering the surface of the younger leafs. Thus, the next analysis focused on trying to compare the relative exploration of each leaf, and the stem apex. It was observed that the apical part of the stem has high and almost constant importance, being explored by most of the bees exploring the resin of *S. parahyba* individual (Figure 5).

**Figure 5:**
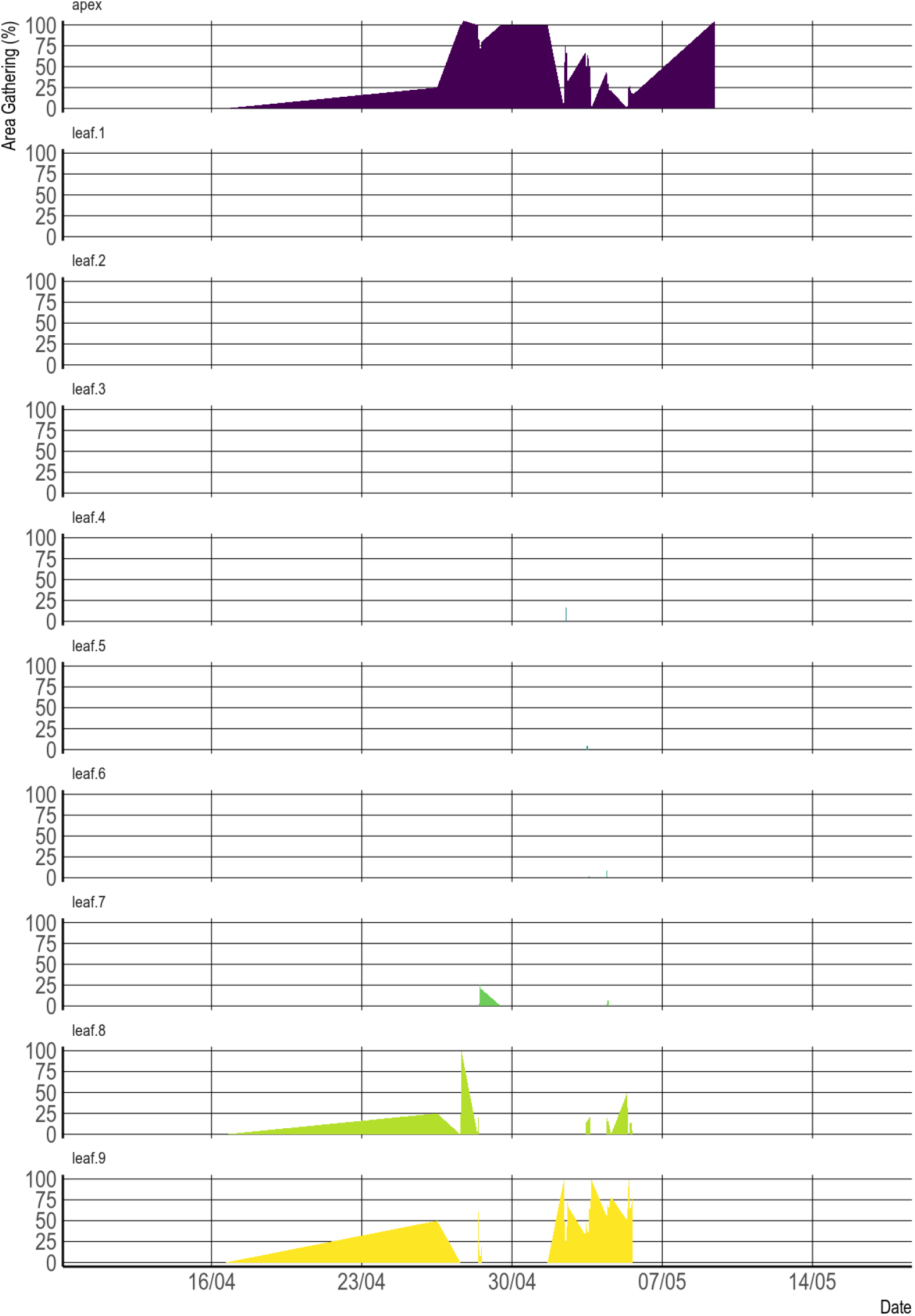
Relative exploration of each leaf from *S. parahyba* individual. Higher values represent more apical leafs.

Comparing the leafs, the most explored are the younger (higher numbers) ones (Figure 5). The alternate phyllotaxy of *S. parahyba* (Lorenzi 2014) result in age differences of each leaf, and this is reflected in a gradual higher use of the youngest ones. It is also possible to follow the growth of importance of the younger leafs, between 16/04 and 25/04.

After one week of rising exploration or young leaves (Figure 5), they suddenly stop exploring them and concentrate their exploration mainly on the stem apex (Figure 5 about 24/04). A few days later, they return to explore the young leaves.

This indicates that the resin deposits are maybe reduced after the intense exploration, thus alternating the exploitation sources make it possible to more effectively obtain the surface resins. This implies that the young part of the plant body is possibly capable of recovering the resin production.

Usually is mentioned that the previous attempts to correlate direct observation of resin gathering and propolis composition were only partially successful because, from several species indicated as resins sources, only some were effectively used to propolis (Bankova, Popova, and Trusheva 2018).

Sundry factors could be responsible for this. First of all, careful observations may be required and correctly documented. In Meliponini, including *T. angustula*, not all the resins were added to propolis, some will be used as resins deposits (piles inside the nest almost made of resins, used as defence) and some will be added to pots (Grüeter 2020). Thus, the proper chemical comparison is required to effectively confirm the use of *S. parahyba* resin as a propolis component.

## Conclusions

*T. angustula* extensively uses the young parts of *S. parahyba* to obtain the surface resin of this plant in most seasons. Resin gathering activity seems to be coherent with hive external activities. Summer had a greater number of bees obtaining resins with a gradual reduction until winter. Temperature and gathering activity was only moderately correlated.

Tree exploration was concentrated on the younger leaves and the stem apex. The alternation between those parts exploration indicates that bees can selectively choose better sources, and possibly plants could restore the surface resin.

Regardless of the restricted reach of this study, which must lead to careful use of the data presented. Proper documentation of the bee behaviour in resin gathering is very important and new. Thus, this study mentioned discoveries aimed to encourage new works in this field.

## References

Akatsu, Ivan Paulo, and Ademilson Espencer Egea Soares. 2009. “Resinas vegetais coletadas por Scaptotrigona (Hymenoptera, Apidae): composição química e atividade atimicrobiana.” PhD thesis, Ribeirão Preto. https://repositorio.usp.br/item/001772961 (https://repositorio.usp.br/item/001772961).

Bankova, Vassya, Davide Bertelli, Renata Borba, Bruno José Conti, Ildenize Barbosa da Silva Cunha, Carolina Danert, Marcos Nogueira Eberlin, et al. 2019. “Standard Methods for Apis Mellifera Propolis Research.” Journal of Apicultural Research 58 (2): 1–49. https://doi.org/10.1080/00218839.2016.1222661 (https://doi.org/10.1080/00218839.2016.1222661).

Bankova, Vassya, Milena Popova, and Boryana Trusheva. 2018. “The Phytochemistry of the Honeybee.” Phytochemistry 155 (November): 1–11. https://doi.org/10.1016/j.phytochem.2018.07.007 (https://doi.org/10.1016/j.phytochem.2018.07.007).

Bisui, Sourabh, Ujjwal Layek, and Prakash Karmakar. 2019. “Comparing the Pollen Forage Pattern of Stingless Bee (Trigona Iridipennis Smith) Between Rural and Semi-Urban Areas of West Bengal, India.” Journal of Asia-Pacific Entomology 22 (3): 714722. https://doi.org/10.1016/j.aspen.2019.05.008 (https://doi.org/10.1016/j.aspen.2019.05.008).

Borges, Francine von B., and Betina Blochtein. 2005. “Atividades externas de Melipona marginata obscurior Moure (Hymenoptera, Apidae), em distintas épocas do ano, em São Francisco de Paula, Rio Grande do Sul, Brasil.” Revista Brasileira de Zoologia 22 (3): 680–86. https://doi.org/10.1590/S0101-81752005000300025 (https://doi.org/10.1590/S0101-81752005000300025).

Campião, Miriam de Souza, Juliana Garlet, and Wescley Viana Evangelista. 2021. “Resistência de moirões de Schizolobium parahyba var. amazonicum tratados pelo método de substituição de seiva ao ataque de cupins.” In, 340–54. https://doi.org/10.37885/210404133 (https://doi.org/10.37885/210404133).

Çelemli, Ömür Gençay. 2013. “Chemical Properties of Propolis Collected by Stingless Bees.” In, edited by Patricia Vit, Silvia R. M. Pedro, and David Roubik, 525–37. New York, NY: Springer New York. https://doi.org/10.1007/978-1-4614-4960-7_39 (https://doi.org/10.1007/978-1-4614-4960-7_39).

Da Silva, Felipe C, Carmen Favaro-Trindade, Severino Matias Alencar, Marcelo Thomazini, and Julio Cesar De Carvalho Balieiro. 2011. “Physicochemical Properties, Antioxidant Activity and Stability of Spray-Dried Propolis.” Journal of ApiProduct and ApiMedical Science 3 (2): 94100. https://doi.org/10.3896/IBRA.4.03.2.05 (https://doi.org/10.3896/IBRA.4.03.2.05).

Dancey, Christine P., and John Reidy. 2020. Statistics Without Maths for Psychology. 8th ed. Harlow, England; New York: Pearson. https://3lib.net/book/17240752/0bdd73 (https://3lib.net/book/17240752/0bdd73).

Divya, K. K., V. S. Amritha, Aswathy Viswanathan, and S. Devanesan. 2020. “Microbial Count in Stingless Bee Honey (Tetragonula Iridipennis (Smith))” 5 (6).

Figueiredo-Mecca, Gláucya de, Luci Rolandi Bego, and Fabio Santos do Nascimento. 2013. “Foraging Behavior of Scaptotrigona Depilis (Hymenoptera, Apidae, Meliponini) and Its Relationship with Temporal and Abiotic Factors.” Sociobiology 60 (3): 267–82. https://doi.org/10.13102/sociobiology.v60i3.267-282 (https://doi.org/10.13102/sociobiology.v60i3.267-282).

Grüeter, Christoph. 2020. “Evolution and Diversity of Stingless Bees.” In, 43–86. Fascinating Life Sciences. Switzerland: Springer Nature. https://doi.org/10.1007/978-3-030-60090-7_2 (https://doi.org/10.1007/978-3-030-60090-7_2).

Kassambara, Alboukadel. 2020. Ggpubr: ‘Ggplot2’ Based Publication Ready Plots. https://CRAN.R-project.org/package=ggpubr (https://CRAN.R-project.org/package=ggpubr).

Kleinert-Giovannini, A., and V. L. Imperatriz-Fonseca. 1986. “Flight Activity and Responses to Climatic Conditions of Two Subspecies of Melipona Marginata Lepeletier (Apidae, Meliponinae).” Journal of Apicultural Research 25 (1): 3–8. https://doi.org/10.1080/00218839.1986.11100685 (https://doi.org/10.1080/00218839.1986.11100685).

Kumazawa, Shigenori, Jun Nakamura, Masayo Murase, Mariko Miyagawa, Mok-Ryeon Ahn, and Shuichi Fukumoto. 2008. “Plant Origin of Okinawan Propolis: Honeybee Behavior Observation and Phytochemical Analysis.” Naturwissenschaften 95 (8): 781. https://doi.org/10.1007/s00114-008-0383-y (https://doi.org/10.1007/s00114-008-0383-y).

Langenheim, Jean H., David E. Lincoln, W. H. Stubblebine, and A. C. Gabrielli. 1982. “Evolutionary Implications of Leaf Resin Pocket Patterns in the Tropical Tree Hymenaea (Caesalpinioideae: Leguminosae).” American Journal of Botany 69 (4): 595–607. https://doi.org/10.1002/j.1537-2197.1982.tb13296.x (https://doi.org/10.1002/j.1537-2197.1982.tb13296.x).

Lorenzi, Harri. 2014. Árvores Brasileiras: manual de identificação e cultivo de plantas arbóreas nativas do Brasil. 5th ed. Vol. 1. Nova Odessa-SP: Plantarum. https://3lib.net/book/3391396/b5a328 (https://3lib.net/book/3391396/b5a328).

Melo, Gabriel A. R. 2020. “Stingless Bees (Meliponini).” In, edited by Christopher K. Starr, 1–18. Cham: Springer International Publishing. https://doi.org/10.1007/978-3-319-90306-4_117-1 (https://doi.org/10.1007/978-3-319-90306-4_117-1).

R Core Team. 2020. R: A Language and Environment for Statistical Computing. Vienna: R Foundation for Statistical Computing. http://www.r-project.org/ (https://www.r-project.org/).

Romão, M.V.V., and V.F. Mansano. 2020. “Schizolobium.” In. Jardim Botânico do Rio de Janeiro. http://reflora.jbrj.gov.br/reflora/floradobrasil/FB23143 (http://reflora.jbrj.gov.br/reflora/floradobrasil/FB23143).

Sawaya, Alexandra Christine Helena Frankland, Juliana Cristina Pereira Calado, Letícia Cavalcanti dos Santoss, Maria Crisstina Marcucci, Ivan Paulo Akatsu, Ademilsoln Espencer Egea Soares, Patrícia Verardi Abdelnur, Ildenize Barbosa da Silva Cunha, and Marcos nogueira Eberlin. 2009. “Composition and Antioxidant Activity of Propolis from Three Species of Scaptotrigona Stingless Bees.” Journal of ApiProduct and ApiMedical Science 1 (2): 3742. https://doi.org/10.3896/IBRA.4.01.2.03 (https://doi.org/10.3896/IBRA.4.01.2.03).

Wickham, Hadley. 2020. Tidy Messy Data: Tidyr. https://CRAN.R-project.org/package=tidyr (https://CRAN.R-project.org/package=tidyr).

